# Comprehensive Model of Cell-to-Cell Cytokinin Transport Reveals A Specific Mode of Cytokinin Riboside Influx

**DOI:** 10.1101/2024.06.04.597342

**Authors:** Daniel Nedvěd, Martin Hudeček, Petr Klíma, Jozef Lacek, Karel Müller, Petr Hošek, Ján Šmeringai, Markéta Pernisová, Václav Motyka, Ondřej Plíhal, Klára Hoyerová

## Abstract

Ribosylated forms of plant hormones cytokinins (CKs) are the dominant CK species translocated at long distances. Their particular roles in plant physiology imply the existence of a yet uncharacterized CK riboside-specific membrane transport system. In this work, we report significant differences in the kinetics of the membrane transport of CK nucleobases and ribosides and the overall affinity of membrane-bound carriers towards the two CK forms. We show that CK ribosides can inhibit the uptake of CK nucleobases in tobacco Bright Yellow 2 cell suspensions but not vice versa, confirming the existence of a membrane transport system that strictly recognizes CK ribosides.

We further characterize the membrane transport of CK nucleobases and ribosides mediated by AtENT3 (EQULIBRATIVE NUCLEOSIDE TRANSPORTER 3), showing its preference towards *trans*-zeatin riboside (tZR) over isopentenyl adenosine (iPR). With the molecular docking and molecular dynamics, we assess the interactions among the side chain of tZR and AtENT3 residues Tyr61 and Asp129, which are conserved in all AtENTs but not in the ENTs from non-plant species. Lastly, we show that *atent3* mutation affects shoot phenotype, demonstrating the impact of CK riboside membrane transport on shoot development.

## 3 Introduction

Cytokinins (CKs) are plant hormones that regulate a great variety of physiological processes, including cell cycle and proliferation (Miller et al., 1956; Schaller et al., 2014), growth and branching of both shoots and roots (Chang et al., 2015; Dello Ioio et al., 2012; Schaller et al., 2014; Skoog and Miller, 1957; Werner et al., 2001), chlorophyll retention and delay of senescence (Dobránszki and Mendler-Drienyovszki, 2014; Richmond and Lang, 1957; Talla et al., 2016) or differentiation of vascular elements (Bishopp et al., 2011; De Rybel et al., 2014; Mähönen et al., 2006).

As signalling molecules, CKs participate in communication between various parts of the plant. They are distributed among tissues and organs through the two vascular pathways - phloem and xylem - but the CK composition in each of them differs, and so presumably do their roles (Corbesier et al., 2003; Hirose et al., 2008; Osugi et al., 2017; Sakakibara, 2021). To reach their eventual destination, CKs have to pass through biological membranes. One possible meaning of membrane transport is simple diffusion, which is described by Fick’s laws (Paul et al., 2014). Due to the hydrophobic character of the inner leaflets of the biological membrane, only small and non-polar molecules can cross the membrane this way. The required characterization applies to CK nucleobases, N^6^-substituted derivatives of adenine, which are the biologically active CK form (Lomin et al., 2015). In contrast, CK ribosides, N^9^*-*ribosylated conjugates of CK nucleobases and the dominant CK components found in the vasculature (Corbesier et al., 2003; Sakakibara, 2021), are bulky and polar, which implies that their diffusion would be inefficient (for more detailed comparison, see <u>Nedvěd et al., 2021</u>). CK nucleobases and ribosides are recognized by membrane-bound carriers, which significantly improves the kinetics of their membrane transport. These carriers belong to the families of PURINE PERMEASES (PUPs) (Hu et al., 2023; Qi and Xiong, 2013; Rong et al., 2024; Xiao et al., 2020, 2019; Zürcher et al., 2016), ATP-BINDING CASSETTES (ABCs) (Jamruszka et al., 2024; Jarzyniak et al., 2021; Kim et al., 2020; Ko et al., 2014; Yang et al., 2022; Zhang et al., 2014; Zhao et al., 2023, 2019), AZA-GUANINE RESISTANT (AZG) (Tessi et al., 2023, 2020), SUGAR WILL EVENTUALLY BE EXPORTED TRANSPORTERS (SWEETs) (Radchuk et al., 2023), and EQUILIBRATIVE NUCLEOSIDE TRANSPORTERS (ENTs) (Girke et al., 2014; Hirose et al., 2008, 2005; Korobova et al., 2021; Sun et al., 2005).

Given that CK ribosides are the main form of CKs transported over long distances, the membrane transport of ribosylated CKs represents a link between the long-distance and cell-to-cell CK distribution. Unlike CK nucleobases, CK ribosides can travel from the root up to the shoot apex and regulate processes such as leaf emergency rate in response to nutrient availability, which likely requires involvement of CK riboside transporters (Davière and Achard, 2017; Landrein et al., 2018; Lopes et al., 2021; Osugi et al., 2017; Sakakibara, 2021).

The physiological importance of CK ribosides implies the existence of a CK riboside-specific system of membrane-bound carriers that are likely separated from the transport of CK nucleobases. Apparent candidates for these carriers are some members of the ENT family mentioned above. AtENT3, 6, 8 from mouse-ear cress (*Arabidopsis thaliana*, L.) (Hirose et al., 2008; Korobova et al., 2021; Sun et al., 2005) and OsENT2 from rice (*Oryza sativa*, L.) (Hirose et al., 2005) have been characterized as CK transporters although only OsENT2 has been shown to directly transport CKs across the biological membrane in a yeast model system.

In this work, we emphasize the importance of CK riboside transport by pin-pointing the different kinetics of CK nucleobase and riboside uptake in the BY-2 cell line (*Nicotiana tabacum*, L. cv Bright Yellow 2), a plant single-cell system (Nagata et al., 1992). Furthermore, we directly monitor AtENT3-mediated CK influx in BY-2, model interactions between AtENT3 and *trans*-zeatin riboside and demonstrate the involvement of AtENT3 in shoot development in *A. thaliana*.

## 4 Material and Methods

### 4.1 Plant Material

We maintained tobacco cell line BY-2 (*N. tabacum* L. cv Bright Yellow 2) in liquid Murashige and Skoog (MS) medium (30 g L^-1^ sucrose, 4.34 g L^-1^ MS salts, 100 mg L^-1^ myo-inositol, 1 mg L^-1^ thiamine, 0.2 mg L^-1^ 2,4-dichlorophenoxyacetic acid, 200 mg L^-1^ KH_2_PO_4_; pH = 5.8), in the dark, at 27 °C, under continuous shaking (150 rpm; orbital diameter 30 mm), and subcultured it every seven days. We cultured the *AtENT3*-expressing transgenic BY-2 cells and calli in the same medium supplemented with 100 mg mL^-1^ cefotaxime and 20 mg mL^-1^ hygromycin.

We grew *A. thaliana* ecotype Columbia 0 (Col-0) and *atent3* T-DNA insertion mutant, obtained from Nottingham Arabidopsis Stock Centre as N631585, on solid MS medium (2.17 g L^-1^ MS salts, 10 g L^-1^ agar; pH = 5.7) in Petri dishes and Klasmann TS-3 fine cultivation substrate (Klasmann-Deilmann GmbH, Germany) in 7.0×7.0×6.5 cm pots. We kept the seeds sown on solid MS medium in the darkness at 4 °C for three days and then cultivated them for eight days under long-day conditions (16 h light/8 h dark) at 20/22 °C in the D-root system (Silva-Navas et al., 2015) using poly klima^®^ climatic growth chambers (poly klima^®^, Germany). We randomly arranged the potted plants of different genotypes in transportable trays with a capacity of 20 pots (4×5 template) and grew them in cultivation chambers – phytotrons (CLF Plant Climatics, Germany) under long-day conditions at 21°C with a LED light intensity of 130 μM m^−2^ s^−1^ and 40-60% relative humidity. Unless stated otherwise, we obtained all chemicals and kits from Sigma–Aldrich Inc.

### 4.2 Transformation of BY-2 Cells

To construct the *XVE::AtENT3* inducible system, we amplified the 1939bp sequence of the *AtENT3* gene from genomic DNA using the forward and reverse *AtENT3* primers with attB1 and attB2 sites, respectively. The primer sequences are listed in Table S1. We cloned the amplified *AtENT3* gene flanked by attB sites into the pDONR207 vector using BP recombination. Subsequently, we transformed the *AtENT3* entry clone into the pMDC7 destination vector (Curtis and Grossniklaus, 2003) by LR recombination. We transformed BY-2 cells by co-cultivation with *Agrobacterium tumefaciens* strain GV2260 (An et al., 1985). We harvested transgenic lines after 4 weeks, cultured them on solid media with kanamycin, and tested for the presence of *AtENT3* via PCR.

### 4.3 Radio-Accumulation Assays

For the radio-accumulation assays, we used BY-2 cell suspensions two days after inoculation. We filtered away the liquid phase of the suspension, twice resuspended the cells in uptake buffer (20 mM 2-morpholin-4-ylethanesulfonic acid, 10 mM sucrose, 0.5 mM CaSO_4_, pH = 5.7), and cultivated them in the dark for 45 and 90 minutes, respectively. The assay itself was initiated by applying a radio-labelled tracer into the cell suspension and terminated after 15-30 minutes. During the assay, we sampled 500 µL of the suspensions in regular intervals. For each sample, we filtered away the liquid phase and treated the cells with 500 µL of 96 % (v/v) ethanol for 30 minutes. Next, we added 4 mL of scintillation cocktail EcoLite(+)^TM^ (MP Biomedicals, CA, USA) to each sample and mixed the samples for 20 min using orbital shaker KS 130 (IKA, Germany) at 480 rpm. The radioactivity in samples was measured using Tri-Carb 2900TR scintillation counter (PerkinElmer, CT, USA).

### 4.4 Mathematical Modelling of Transport Kinetics

To describe the kinetics of the CK membrane transport in BY-2 cell culture, we adapted the model published by Hošek et al. (2012). We introduced first-order rate constants *I* and *E* to characterize the influx and efflux of a radio-labelled tracer, respectively. To account for the tracer adsorption to the cell surfaces, we included a factor *K*. To estimate the values of *I*, *E*, and *K*, we fitted experimental data from radio-accumulation assays with equation:

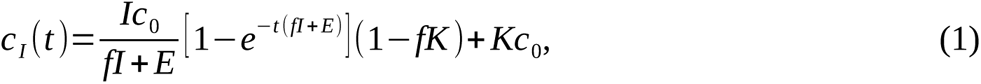

where *t* and *c_I_* are matrices of time points and measured intracellular concentrations, respectively (with each row composed of data points from one assay and different rows representing different assays), *f* is a factor correcting different sizes of the intra- and extracellular spaces, and *c_0_* is the initial extracellular concentration of the tracer. When comparing the effects of the *AtENT3* expression or a chemical treatment on the tracer influx, we constrained the model to keep common values of *E* and *K* for all assays in the dataset. For assays involving chemical treatment during the tracer accumulation (as opposed to the treatment before the tracer addition), we used an expanded form of equation (1):

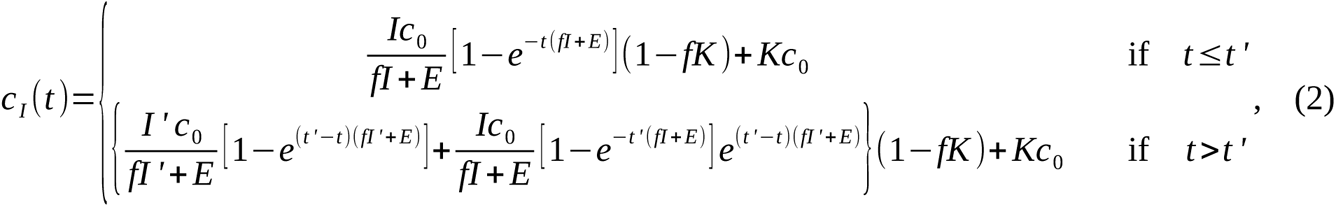

where *t ’* is the treatment time and *I ’* is the influx rate constant after the treatment. For step-by-step derivations of equations (1) and (2), see the Supplementary methods.

To evaluate the affinity of the membrane transport system towards a tracer or the inhibition effect of a competitor, we adapted a saturation model published by Delbarre et al. (1996):

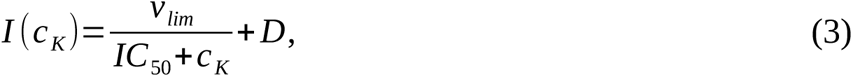

where *c_K_*is the concentration of a competitor (either the non-labelled counterpart of the tracer or another chemical substance), *v_lim_* is the limit transport rate, *IC_50_* is a *c_K_* value for which the transport rate equals to half of *v_lim_*, and *D* is the rate constant of the influx that remains even when the transport system is fully saturated.

All fits were performed using the "curve_fit" method of the SciPy Python library (Virtanen et al., 2020) with arguments "ftol=1e-15" and "xtol=1e-15". The initial guesses were 10^-3^ for *I* and *E*, 0 for *K*, and 1 for *v_lim_*, *IC_50_* and *D*. All parameters were restricted to be non-negative. To visualize tracer accumulation in the cells, we used equation (1) with the optimized values of *I* and *E,* while setting *K* to 0.

### 4.5 Molecular Docking

For molecular docking, we downloaded AlphaFold-predicted structural models (Jumper et al., 2021) from the AlphaFold Protein Structure Database (https://alphafold.ebi.ac.uk/). We prepared the protein and ligand files and performed the docking procedure using the AutoDockFR (ADFR) software suite (Ravindranath et al., 2015; Zhao et al., 2006). The ligands were initially placed in the central cavity of the protein. The centre and dimension of the affinity grids were determined automatically by the ADFR program "agfr". Each docking consisted of 50 runs and each run performed 50 million evaluations. Residues Gln133, Arg312, Leu397, and Asp129 of AtENT3 were set flexible. To visualize the protein-ligand structures, we used ChimeraX (Goddard et al., 2018), PyMol (Schrödinger, LLC, 2015), and LigPlot+ software (Laskowski and Swindells, 2011). We used MAFFT with the automatic algorithm selection (Katoh and Standley, 2013) to align the ENT sequences and JalView for alignment visualization (Waterhouse et al., 2009).

### 4.6 Molecular Dynamics

For the molecular dynamic simulations, we built a rhombic dodecahedron-shaped simulation box consisting of the protein-ligand complex in 150 mM aqueous NaCl solution and additional ions to neutralize the electric charge. To prevent the complex from interacting with its own image, we set its distance from the box edges to 1.0 nm. We ran energy minimization using the steepest descent algorithm, a 100 ps-long simulation under the *NVT* ensemble (constant particle amount, volume, and temperature), a 100 ps-long simulation under the *NPT* ensemble (constant particle amount, pressure, and temperature), and finally 100 or 200 ns-long unbiased simulation. In the *NVT* and *NPT* runs, we applied constraints with force constants of 1000 kJ mol^-1^ nm^2^ to all non-hydrogen atoms. For all runs, we used the CHARMM36 force field (Best et al., 2012). For simulation parameters, see Table S2. We used the GROMACS software suite (Abraham et al., 2024, 2015; Páll et al., 2015) to parametrize the protein, build the simulation box, carry out the simulations, perform the cluster analysis, and calculate the distributions of distances and angles over the trajectories. Through GROMACS, we also used particle mesh Ewald method to evaluate long-range interactions (Essmann et al., 1995), LINCS algorithm to solve constraints (Hess, 2008), and SETTLE algorithm to treat water molecules (Miyamoto and Kollman, 1992). To parametrize the ligand, we used CGenFF program (Vanommeslaeghe et al., 2012; Vanommeslaeghe and MacKerell, 2012). To calculate fractional occupancies of the system by water molecules, we used VMD software (Humphrey et al., 1996).

### 4.7 Plant Phenotyping and Imaging

For phenotyping of 8 day-old *A. thaliana* plants grown on agar, we isolated their shoots, placed these shoots on Petri dishes and scanned them from top view using Epson Perfection V700 Photo (Epson, Japan). To image the potted plants, we used the PlantScreen^TM^ Compact System (PSI, Czechia) equipped with PSI DUAL camera containing two 12.36-megapixel complementary metal-oxide-semiconductor (CMOS) sensors: a colour Sony IMX253LQR-C sensor for RGB structural imaging and a monochromatic Sony IMX253LLR-C for chlorophyll fluorescence measurement (Sony, Japan). For the fluorescence measurement, we used Quenching analysis protocol. Raw data were automatically processed using the PlantScreen^TM^ Analyzer software (PSI). The imaging was performed according to a previously published protocol (Šmeringai et al., 2023).

### 4.8 Image Processing

To process images of agar-grown plants, we transformed the images of isolated plant shoots from the RGB to L*a*b* space and segmented the shoots by applying the thresholds *a\** <= -9.5, *b\** >= - 9.5, and *L\** >= 18.5, based on estimates obtained by multi-Otsu method (Liao et al., 2001). In the binary mask, we removed all objects smaller than 2048 pixels, performed morphological closing using a disk-shaped footprint with a radius of 8 pixels, and then removed all objects smaller than 8192 pixels. Finally, we measured the areas of all remaining objects in the image. To implement the techniques listed above, we used Python scikit-image library (van der Walt et al., 2014). We processed images of potted plants according to Šmeringai et al. <u>(2023)</u>. For miscellaneous image manipulations, we used the GNU Image Manipulation Program (The GIMP Team, 2024).

### 4.9 Reverse Transcription Quantitative PCR

We isolated total RNA isolated from plant shoots using the RNeasy Plant Mini kit (Qiagen, Germany) and treated with DNA-Free kit (Thermo Fischer Scientific, MA, USA). We evaluated the purity, concentration, and integrity of RNA on 0.8% agarose gels (v/w) and by the RNA Nano 6000 Assay Kit using Bioanalyzer instrument (Agilent Technologies, CA, USA). For reverse transcription of approximately 1 mg of the DNAse-treated RNA, we used M-MLV Reverse Transcriptase, RNase H(-), Point Mutant (Promega, WI, USA). We performed the quantitative PCR using GoTaq qPCR Master Mix (Promega, WI, USA) at the annealing temperature of 58 °C on LightCycler480 instrument (Roche, Switzerland). PCR efficiency was estimated using serial dilution of template cDNA. We calculated the relative expression level, *REL*, as follows:

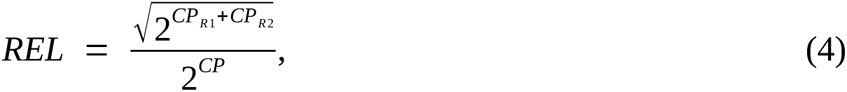

where *CP*_*R*1_ and *CP*_*R*2_ are the crossing points of the reference gene 1 and 2, respectively, and *CP* is the crossing point of the target gene. We used *A. thaliana* elongation factor 1a (AtEF1a) and actin 2 (AtACT2) as reference genes. We verified positive transcript levels and the quality of PCR by the melting curve analysis. The primer sequences are listed in Table S1.

## 5 Results

### 5.1 Transport of Cytokinin Ribosides in BY-2 Cells Occurs with Kinetics Distinct from Cytokinin Nucleobases and Depends on Proton Gradient

To find out whether the membrane transport kinetics of ribosylated CKs differ from the kinetics of CK nucleobases, the biologically active CK form, we measured the uptake of various radio-labelled CK tracers in tobacco Bright Yellow 2 (BY-2) cell cultures (Nagata et al., 1992), a model plant single-cell system. The radio-labelled tracers comprised four CK nucleobases: [3H]-*trans*-zeatin (tZ), [3H]-dihydrozeatin (DHZ), [3H]-isopentenyl adenine (iP), [3H]-benzyladenine (BA), and four CK ribosides: [3H]-*trans*-zeatin riboside (tZR), [3H]-dihydrozeatin riboside (DHZR), [3H]-isopentenyl adenosine (iPR), and [3H]-benzyladenosine (BAR). With this selection, we also included CKs with diverse characters of their side chains (N^9^-bound moieties), namely those with unsaturated hydroxylated chains (tZ, tZR), saturated hydroxylated chains (DHZ, DHZR), unsaturated aliphatic chains (iP, iPR), and aromatic chains (BA, BAR).

To obtain comparable kinetic parameters for each tracer, we fitted the transport model given by equation (1) into the dataset of sampling time points and measured the radioactivities corresponding to the intracellular concentrations of accumulated CK tracers. We used the influx rate obtained by modelling, *I*, to compare the uptake kinetics of different CK tracers. Comparing the median values of *I* obtained for tZ (17.86×10^-3^ s^-1^), DHZ (14.94×10^-3^ s^-1^), and iP (11.58×10^-3^ s^-1^) with the median *I* values obtained for tZR (3.45×10^-3^ s^-1^), DHZR (2.38×10^-3^ s^-1^), and iPR (5.44×10^-3^ s^-1^) shows that BY-2 cells accumulate CK nucleobases with non-aromatic side chains more readily than their respective ribosides. Regarding the side chain composition, the differences in transport among tZ-type, DHZ-type, and iP-type CKs are less pronounced than the differences between CK nucleobases and ribosides. The median values of *I* obtained for BA (9.14×10^-3^ s^-1^) and BAR (16.22×10^-3^) show a difference between the uptake of the nucleobase and the riboside as well, but in this case, the more readily transported form is the riboside. The distributions of *I* values and the accumulation trends modelled for each assay are depicted in Figure 1. For the complete list of kinetic parameters and their statistical analysis, see Table S3 and Table S4, respectively.

**Figure 1:**
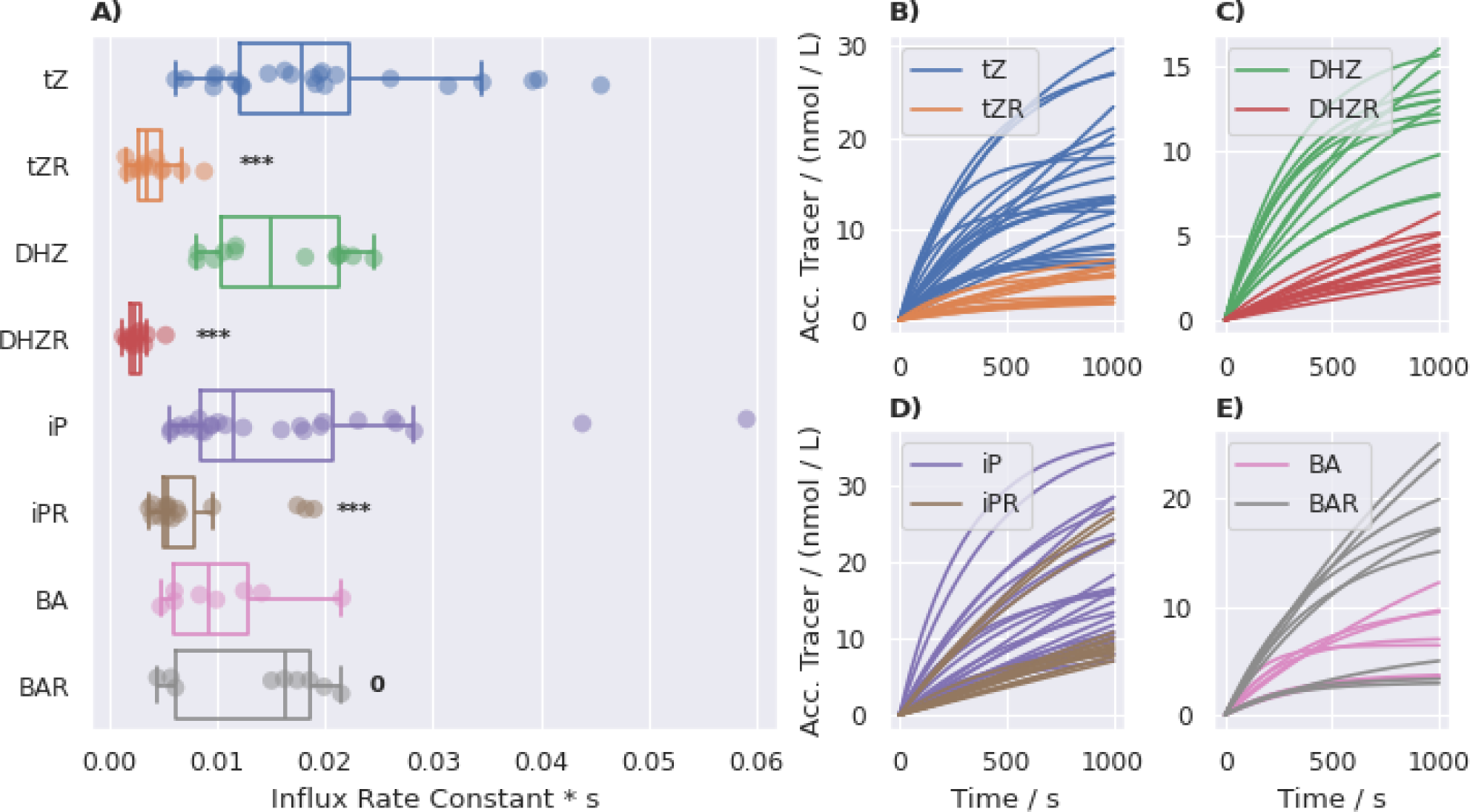
Characterization of the CK membrane transport in tobacco BY-2 cells. **A**: Estimated values of the influx rate constant (*I*) for different radio-labelled CK tracers obtained by fitting equation (1) into data from radio-accumulation assays. **B-E**: Comparison of the accumulation trends (concentration of the accumulated tracer over time) between CK nucleobases and their ribosylated forms. The curves are aligned by setting *K* = 0 and *c_0_* = 2 nM for each assay. *P*-values obtained from the one-way ANOVA test comparing *I* values for the corresponding pairs of CK nucleobases and ribosides: 0 (*P* > 0.1), * (0.1 >= *P* > 0.05), ** (0.05 >= *P* > 0.01), *** (*P* <= 0.01). Acc.: accumulated.

To confirm that the observed uptake of CK nucleobases and ribosides occurs by carrier-mediated transport, we performed a series of assays in which we accumulated [3H]-tZ, [3H]-tZR, [3H]-iP or [3H]-iPR together with their non-labelled counterparts (so-called competitors) in concentrations of 0, 2 nM, 20 nM, 200 nM, 2 µM, and 20 µM. Each dataset, consisting of experiments performed with one tracer and all concentrations of the corresponding non-labelled competitor, was fitted with the constrained variant of equation (1), i.e. with single values of *E* and *K* for the whole dataset. The uptake of CK nucleobases and ribosides is subject to dose-dependent inhibition by their non-labelled variant, as expected of the membrane-bound carriers becoming saturated (Figure 2A-D). To evaluate this inhibition effect numerically, we fitted the *I* parameter values obtained from the assays with competitors using the saturation model given by equation (3) (Delbarre et al., 1996). The estimated *IC_50_* values obtained for tZ (112.21 nM), tZR (2.33 µM), iP (27.25 nM), and iPR (2.65 µM) show that CK nucleobases are transported with slightly higher affinity than the corresponding ribosides, which further indicates the distinct transport properties of the two CK forms (Figure 2E-H). For the complete list of kinetic parameters, see Table S5.

**Figure 2:**
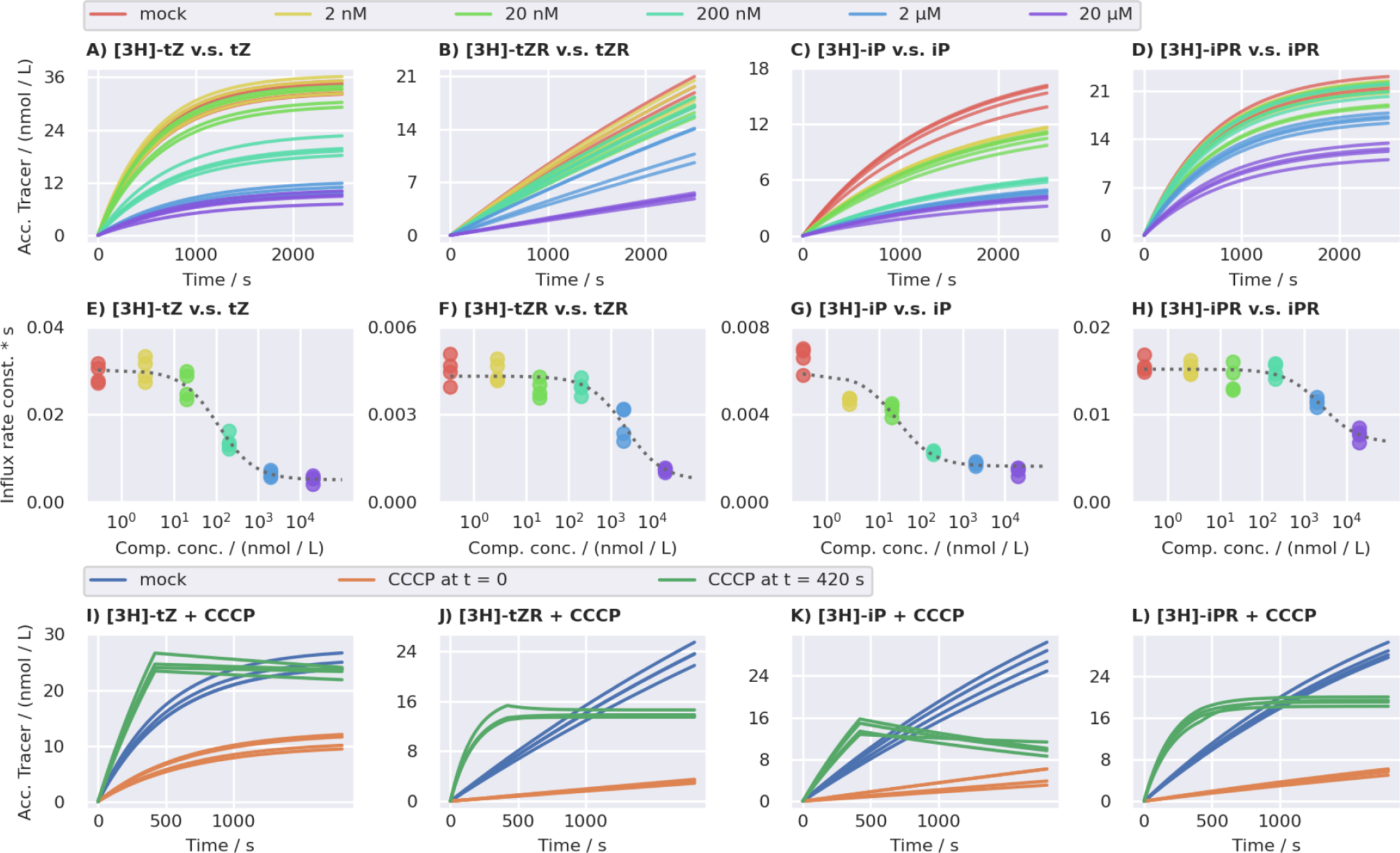
Saturation of the CK membrane transport in tobacco BY-2 cells. **A**-**D**: Accumulation trends (concentration of the accumulated tracer in time) of radio-labelled CK in BY2 cell inhibited by increasing concentrations of their non-labelled counterparts. The shape of the curves is determined by the *I* and *E* values obtained by fitting equation (1) into data from radio-accumulation assays. For the visualization purposes, *K* is set to 0 and *c_0_* to 2 nM for each assay. **E-H**: Dependence of *I* values obtained from the mathematical modelling of radio-accumulation data on the concentration of the non-labelled competitors. The plotted data points are further fitted with equation (3) to obtain the saturation parameters. The fit of equation (3) is represented by grey dashed curves. The *I* values correspond to the curves depicted in A-D. **I-L**: Accumulation trends of radio-labelled CK tracers in presence of 50 µM carbonyl cyanide 3-chlorophenylhydrazone (CCCP). Acc.: accumulated.

To assess the thermodynamic aspect of the uptake of tZ, tZR, iP, and iPR in the BY-2 cells, we performed a series of accumulation assays in suspensions treated with 50 µM protonophore carbonyl cyanide 3-chlorophenylhydrazone (CCCP) in dimethyl sulfoxide (DMSO). CCCP uncouples electron transfer from oxidative phosphorylation (Cavari et al., 1967; Cunarro and Weiner, 1975; Heytler, 1963), thus inhibiting proton gradient-dependent transport processes (Alexander et al., 2018; Culos and Watanabe, 1983; Stoffer-Bittner et al., 2018). We performed the same assays using cells treated with the corresponding amount of DMSO alone (mock treatment) as control. We fitted all data with equation (1) to obtain *I* values. The CCCP treatment decreased the medians of *I* (in comparison with the mock treatment) for all four tracers: from 24.24×10^-3^ to 8.89×10^-3^ s^-1^ for tZ, from 7.78×10^-3^ to 0.91×10^-3^ s^-1^ for tZR, from 9.89×10^-3^ to 1.46×10^-3^ for iP, and from 11.33×10^-3^ to 1.86×10^-3^ s^-1^ for iPR (Figure 2I-L), indicating that the uptake of both CK nucleobases and ribosides at least partially occurs in a proton gradient-dependent manner. For the complete list of kinetic parameters and their statistical analysis, see Table S6 and Table S7, respectively. To observe the immediate response of the CK influx to the uncoupling of the proton gradient, we performed another set of assays in which we treated the cells with 50 µM CCCP in DMSO seven minutes after the onset of the accumulation (i.e. after adding the tracer). We fitted the measured data with equation (2) to visualize the response (for the estimated kinetic parameters, see Table S8). The fits show that after the treatment, the intracellular concentrations of CK nucleobases start to decrease, while the concentrations of CK ribosides stop increasing and remain constant (Figure 2I-L). This trend could indicate the presence of CCCP-resistant exporters of CK nucleobases.

### 5.2 BY-2 Cells Possess a Riboside-Specific Transport System Not Recognizing Cytokinin Nucleobases as Substrates

The different affinities of the BY-2 membrane-bound carriers towards CK nucleobases and ribosides (Figure 2E-H) imply either that both CK types are recognized by the same set of carriers (with the nucleobases being slightly preferred) or that there are two sets of carrier, one for nucleobases and one for ribosides, that function independently of one another. To see which of these models characterizes the CK transport in BY-2 cells better, we paired nucleobase tracers with riboside competitors and vice versa (i.e. [3H]-tZ with tZR, [3H]-tZR with tZ, [3H]-iP with iPR, and [3H]-iPR with iP) and repeated the radio-accumulation assays with increasing concentrations of non-labelled CKs. We fitted the experimental data with equation (1) to assess kinetic parameters for each assay (Figure 3A-D) and the obtained values of *I* with equation (3) to estimate the *IC_50_* for each competitor (Figure 3E-H). The estimated *IC_50_* values are 18.77 µM ([3H]-tZ vs tZR), 90.82 µM ([3H]-tZR vs tZ), and 1.00 µM ([3H]-iP vs iPR). No saturation occurs for the [3H]-iPR vs iP variant (i.e. *IC_50_* diverges towards infinity). For the complete list of kinetic parameters, see Table S5. These results show that the inhibition of the CK nucleobase uptake by CK ribosides is significantly weaker than the inhibition by non-labelled nucleobases and vice versa, supporting the model of two independent carrier sets.

**Figure 3:**
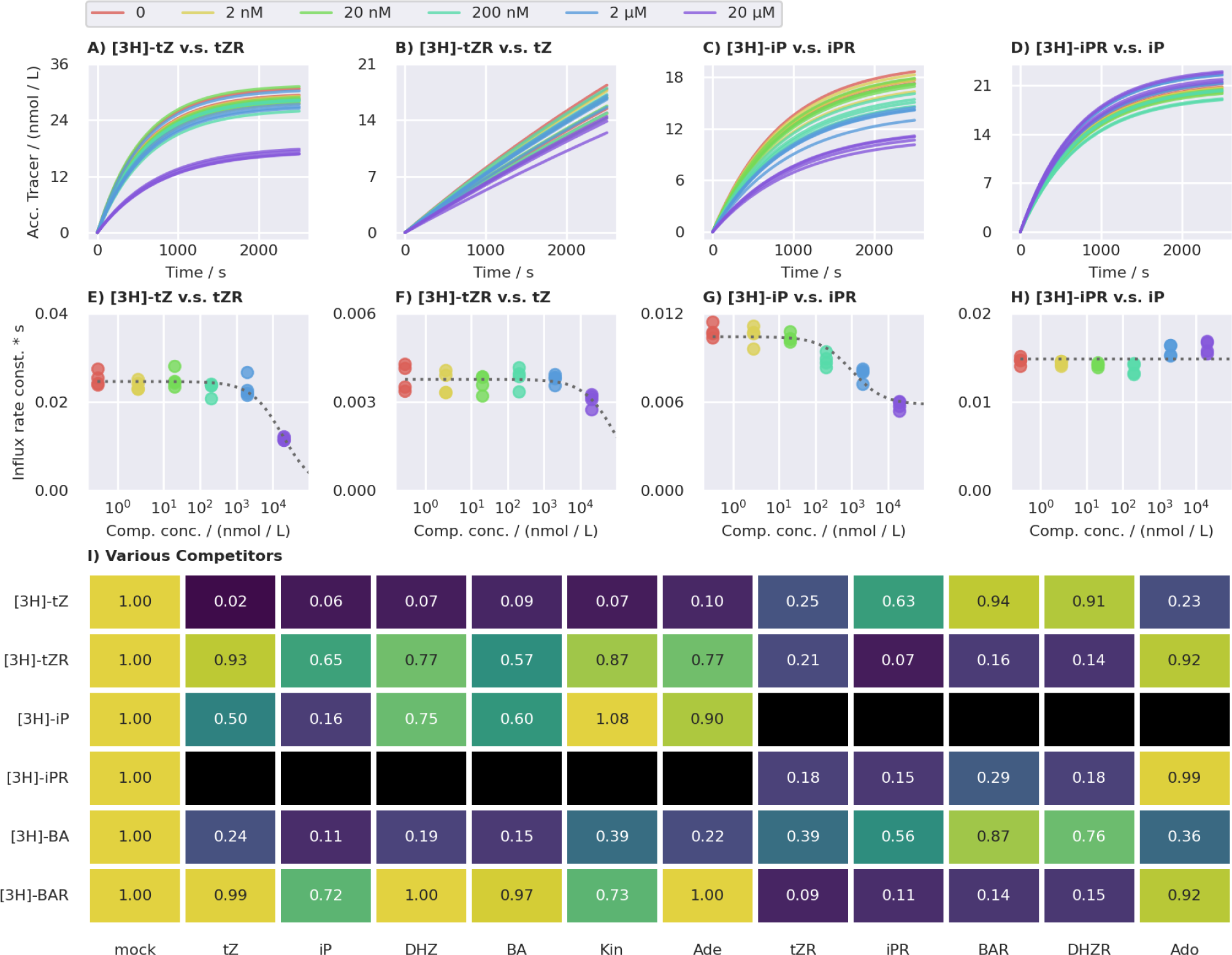
Substrate specificity of CK membrane-bound transport systems. **A**-**D**: Accumulation trends (concentration of the accumulated tracer over time) of radio-labelled CK in BY-2 cell sinhibited by increasing concentrations of chemically diverse non-labelled substances. The shape of the curves is determined by the *I* and *E* values obtained by fitting equation (1) into data from radio-accumulation assays. For the visualization purposes, *K* is set to 0 and *c_0_* to 2 nM for each assay. **E-H**: Dependence of the *I* values obtained from the mathematical modelling of radio-accumulation data on the concentration of the non-labelled competitors. The plotted data points are further fitted with equation (2) to obtain the saturation parameters. The fit of equation (2) is represented by grey dashed curves. The *I* values correspond to the curves depicted in A-D. **I**: Fold changes of the influx rate constants estimated for various combinations of radio-labelled CK tracers and 20 µM non-labelled competitors. Black cells denote non-tested combinations. Acc.: accumulated.

To confirm this trend, we tested the inhibitory effect of more CK- and adenine-based competitors (adenine, tZ, iP, BA, DHZ, kinetin, adenosine, tZR, iPR, BAR, and DHZR) on the uptake of [3H]-tZ, [3H]-tZR, [3H]-iP, [3H]-iPR, [3H]-BA, and [3H]-BAR. The concentration of all competitors was 20 µM. We fitted the measured data with the constrained variant of equation (1) and compared the median values of *I*. The uptake of [3H]-tZ and [3H]-BA decreases (two to four times) in the presence of all tested nucleobases, as well as tZR, adenosine, and iPR ([3H]-BAR only), whereas other tested ribosides cause mild to none uptake inhibition. The uptake of [3H]-iP is about three times reduced by its non-labelled variant but only mildly reduced by other tested compounds. The uptake of all three labelled ribosides is efficiently inhibited by all tested ribosides with a striking exception of adenosine and only mildly inhibited by tested nucleobases (Figure 3I; for the estimated kinetic parameters, see Table S9). The results presented so far indicate the existence of at least two systems mediating the CK membrane transport - one can recognize both CK nucleosides and ribosides (with a slight preference towards the former), while the other is strictly riboside-specific.

### 5.3 AtENT3 Transport Cytokinin Nucleobases and Ribosides, Preferring *trans*-Zeatin Riboside over Isopentenyl Adenosine

To see how our previous conclusions about the CK membrane transport as a whole apply to individual membrane-bound carriers, we decided to proceed with the expression of a previously characterized transporter of CK ribosides in BY-2 cells and measure its contribution to the uptake of tZ, iP, tZR, and iPR to assess its specificity. We have focused on members of the ENT family, as some of them have been linked to the transport of CK ribosides (Girke et al., 2014).

To determine whether tobacco ENTs could be responsible for the CK uptake in BY-2 cells, we searched for expression of *ENT* genes in a previously published BY-2 transcriptome (Müller et al., 2021). All tobacco ENTs listed in the UniProtKB database (The UniProt Consortium, 2023) are homologs of AtENT1, 3 or 8. Of these, only the homologs of *AtENT1* and *3* genes are expressed in BY-2 (Figure 4A; for numerical values, see Table S10), implying that measurements performed on AtENT1 or 3 may reflect the transport trends described above. We eventually decided to further work with AtENT3, given the previous reports on its effects on the CK homeostasis and plant sensitivity to exogenously applied CKs (Korobova et al., 2021; Sun et al., 2005).

**Figure 4:**
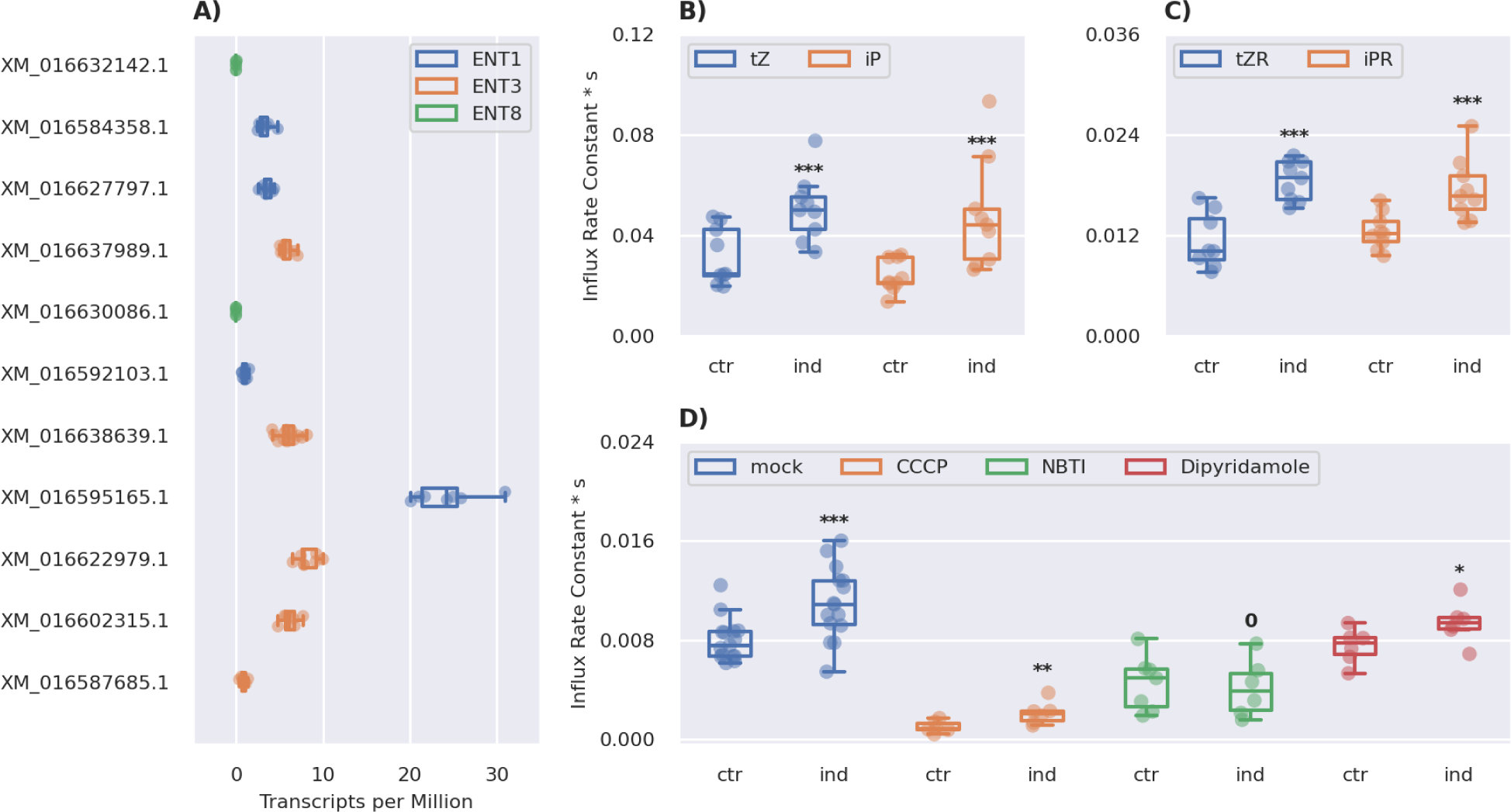
The effect of *AtENT3* expression on the CK uptake in tobacco BY-2 cells. **A**: Expression of *AtENT* homologues in two day-old BY-2 cultures. The identifiers on the vertical axis correspond to accessions in the NCBI (National Center for Biotechnology Information) Gene database (accessed on 17 April 2024). Data were obtained through the GEO (Gene Expression Omnibus) database under the accession of GSE160438 (Müller et al., 2021). **B-C**: Optimized values of the influx rate constant, *I*, obtained by fitting equation (1) into data from radio-accumulation assays measuring the uptake of radio-labelled CK nucleobases and ribosides in the *AtENT3*-harbouring BY-2 cells under an estradiol-inducible promoter without (ctr) or with the induction (ind) of *AtENT3* expression. **D**: Optimized values of *I* for the uptake of radio-labelled tZR in the *AtENT3*-harbouring BY-2 cells without or with the induction of *AtENT3* expression and without (mock) or with transport inhibitors *S*-(4-nitrobenzyl)-6-thioinosine (NBTI), dipyridamole (DiPy), and carbonyl cyanide 3-chlorophenylhydrazone (CCCP). All inhibitors were applied at a concentration of 10 µM. *P*-values obtained from the one-way ANOVA test comparing *I* values for induced and control cell lines: 0 (*P* > 0.1), * (0.1 >= *P* > 0.05), ** (0.05 >= *P* > 0.01), *** (*P* <= 0.01).

To directly monitor the transport activity of AtENT3 towards CKs, we introduced the estradiol-inducible *XVE*::*AtENT3* gene construct to BY-2 cells. Using these transformed cells, we performed radio-accumulation assays with [3H]-tZ, [3H]-tZR, [3H]-iP, and [3H]-iPR as tracers. We performed each assay in non-induced (control) and induced cells to assess the contribution of AtENT3 to the overall CK uptake. The induced cells were treated with 1 µM estradiol in DMSO and the control cells with the corresponding amount of DMSO. The measured data were fitted with the constrained variant of equation (1) to estimate *I* values. The medians of *I* increased for all four tracers: from 24.56×10^-3^ to 50.03×10^-3^ s^-1^ for tZ, from 10.16×10^-3^ to 18.82×10^-3^ s^-1^ for tZR, from 21.34×10^-3^ to 44.27×10^-3^ s^-1^ for iP, and from 12.19×10^-3^ to 16.69×10^-3^ for iPR (Figure 4B-C). For the complete list of kinetic parameters and their statistical analysis, see Table S11 and Table S12, respectively. These results show that AtENT3 transports nucleobases and ribosides, implying that AtENT3 is likely not a part of the previously described CK riboside-specific system and that strict carriers of ribosylated CKs remain to be identified. Focusing on the results obtained for the accumulation of tZR and iPR, we saw that AtENT3 boosts the influx rate of the former more than the latter. The influx rates of tZ and iP are boosted similarly, suggesting that CK nucleobases are transported with a different mechanism, which does not allow discrimination based on the character of the CK N^6^-bound side chain.

To further confirm that AtENT3 is responsible for the increase in CK riboside uptake in the induced cells, we examined how AtENT3-mediated uptake of tZR changes after application of CCCP and two inhibitors of nucleoside uptake, *S*-(4-nitrobenzyl)-6-thioinosine (NBTI) (Karbanova et al., 2020; Ward et al., 2000; Wright and Lee, 2019) and dipyridamole (DiPy) (Newell et al., 1986; Woffendin and Plagemann, 1987). NBTI and DiPy inhibit the uptake of adenosine by AtENTs (Li et al., 2003; Möhlmann et al., 2001; Wormit et al., 2004). To assess the effects of the inhibitors, we performed accumulation assays with [3H]-tZR as a substrate, in non-induced (control) or induced cell lines and with or without CCCP, NBTI or DiPy. All inhibitor we dissolved in DMSO and used at the concentration of 10 µM. For mock treatment, we used the corresponding amount of DMSO alone. We fitted the measured data with the constrained variant of equation (1). In mock-treated cells, the uptake of tZR significantly increases (from 7.56×10^-3^ to 10.87×10^-3^ s^-1^) due to the induction of *AtENT3* expression. In CCCP-treated cells, the overall tZR uptake drops, but there is still a difference between control and induced cells (the median of *I* increases from 0.85×10^-3^ to 2.05×10^-3^ s^-1^). In NBTI-treated cells, there is no significant difference between the control and induced cells, indicating strong inhibition of AtENT3 by NBTI. Finally, in DiPy-treated cells, the median of *I* mildly increases from 7.73×10^-3^ to 9.39×10^-3^ s^-1^, suggesting partial inhibition of AtENT3 (Figure 4D). For the complete list of kinetic parameters and their statistical analysis, see Table S13 and Table S14, respectively. The results of the competition assays show that AtENT3 is inhibited by NBTI and (to a lesser extent) DiPy, two typical inhibitors of adenosine uptake. However, it is resistant to CCCP, suggesting that AtENT3 mediates facilitated diffusion rather than active transport. The resistance of AtENT3 towards CCCP could also be related to the milder response of CK riboside uptake to the CCCP treatment (compared to CK nucleobases) in wild-type BY-2 cells (Figure 2I-L).

### 5.4 A Computational Approach Reveals A Non-Conserved tZR-Interacting Motif in AtENT Sequences

To assess the molecular interactions responsible for CK binding to AtENT3, we performed molecular docking of tZR into a predicted structure of AtENT3 obtained with AlphaFold (Jumper et al., 2021). The best-docked pose of tZR is located in a central cavity outlined by transmembrane helices (TMs) 1, 2, 4, 5, 7, 8, 10, and 11. The ENT3 residues interacting with the docked pose of tZR comprise Leu31, Trp34, Asn35, Tyr61, Gln62, Asp129, Gln133, Tyr272, Leu276, Tyr304, Asn305, Asp308, Lys312, Asn365, Leu396, Leu397, and Ile400 (Figure 5A). This pose roughly corresponds to the sites occupied by the adenosyl moiety of NBTI in human HsENT1 (Wright and Lee, 2019) (PDB code: 6OB6) and by inosine in PfENT1 from the parasite *Plasmodium falciparum* (Wang et al., 2023) (PDB code: 7WN1; see Figure 5C). Moreover, the CK-interacting residues Leu31, Trp34, Gln133, Tyr304, Asp308, and Lys312 correspond to residues reported to bind respective ligands in 6OB6 and 7WN1. We also performed docking of iPR, tZ, and iP. For all best-docked poses, see Figure S1-4.

**Figure 5:**
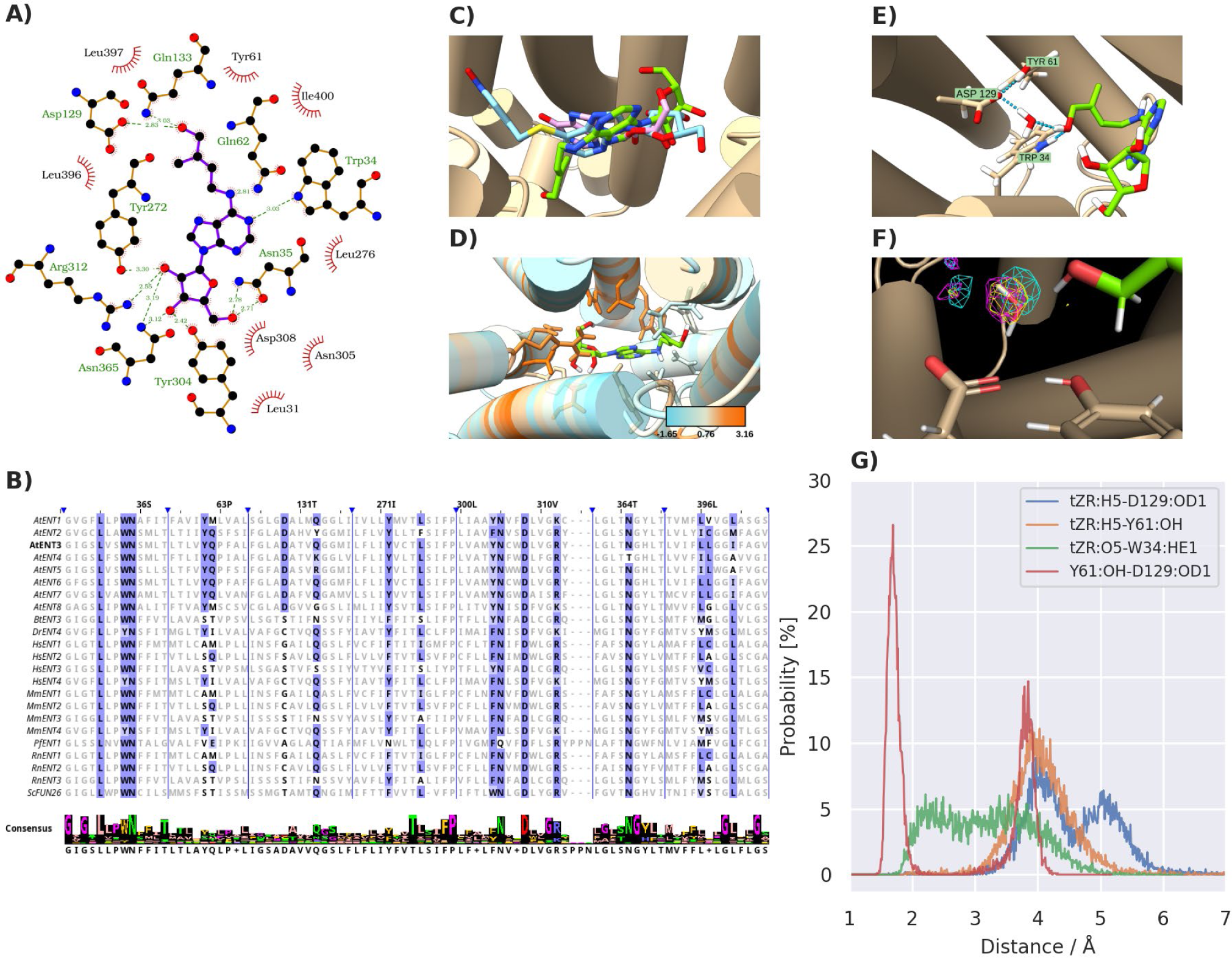
Computational assessment of the interactions between the AlphaFold-predicted structure of AtENT3 and tZR. **A**: Schematic representation of the best docked pose of tZR in the binding cavity of AtENT3. Green dashed lines represent hydrogen bonds with lengths given in Å. Short red rays represent hydrophobic interactions. Visualized in LigPlot+. **B**: Sequential alignment of plant, animal, parasitic, and yeast ENTs. The residues of AtENT3 interacting with the docked pose of tZR and their homologs are shown in bold. Their conservation among the presented species is depicted by the differential intensity of the blue highlight. Blue vertical lines mark breaks in the sequences. The labels in the header of the alignment denote the residues of AtENT3 found at the given position. The consensus sequences and the logotype of the alignment segments are given at the bottom. Visualized in Jalview. **C**: Superimposition of AtENT3 (tan cylinders) with the docked pose of tZR (green) and the experimental poses of NBTI (light blue) in HsENT1 (PDB code: 6OB6) and inosine (pink) in PfENT1 (PDB code: 7WN1). The amino acid residues of 6OB6 and 7WN1 are hidden. **D**: Sequence conservation of AtENT3 residues interacting with the docked pose of tZR calculated by the AL2CO program (Pei and Grishin, 2001) from the alignment depicted in B. Larger numbers indicate higher conservation. **E**: Hydrogen bonding among tZR, Trp34, Tyr61 and Asp129, and a water molecule in the system equilibrated by molecular dynamics (MD). Hydrogen bonds are depicted as light blue dashed lines. Images C-E are visualized in UCSF ChimeraX. **F**: Fractional occupancies of the MD simulation grid by water molecules. The meshes represent isosurfaces with the fractional occupancy of 90%. Differently coloured meshes correspond to three independent MD simulation runs. Visualized in PyMol. **G**: Representative distributions of atomic distances involving the hydrogen (H5) and oxygen (O5) of the side-chain carboxyl of tZR, a carboxylic oxygen of Asp129 (OD1), the phenolic oxygen of Tyr61 (OH), and the nitrogen-bound hydrogen of Trp34 (HE1).

To assess the conservation of the CK-binding residues among the known members of the ENT family, we aligned sequences of the reviewed ENT proteins present in the UniProtKB database (The UniProt Consortium, 2023). These proteins are AtENT1-8, BtENT3 from cattle, DrENT4 from zebrafish, HsENT1-4, MmENT1-4 from mouse, PfENT1, RnENT1-3 from rat, and ScFUN26 from yeast. The alignment shows that the residues interacting with the ribosyl moiety of tZR are generally more conserved than those interacting with the heterocycle and the side chain (Figure 5B, D), suggesting that the binding cavities of different ENTs are all shaped to recognize nucleosides but with different specificities towards various aglycones.

To estimate the stability of predicted AtENT3-tZR interactions, we performed molecular dynamic simulations with a system consisting of the AtENT3-tZR complex in water and 150 mM NaCl. To equilibrate the system, we ran a single 200 ns-long simulation. Through cluster analysis of the 200 ns-long trajectory, we obtained a representative system conformation (corresponding to the frame at t = 177.72 ns). In this conformation, we observed interactions between the side-chain hydroxyl group of tZR and residues Trp34, Tyr61, and Asp129, mediated by a water molecule (Figure 5E). Tyr61 and Asp129 are conserved among AtENTs but not among ENTs from other species listed in Figure 5B, suggesting they might have a unique role in binding CK substrates.

Next, we ran three parallel 100 ns-long simulations, starting from the system conformation obtained through the cluster analysis. To confirm that a water bridge contributes to the stabilization the side-chain hydroxyl of tZR, we calculated the fractional occupancies of water molecules in the system during the simulations. In the space surrounded by the side-chain hydroxyl of tZR and the side chains of Tyr61 and Asp129, the fractional occupancy of water reaches a local maximum of approximately 90%, indicating that this space is occupied by water for the most simulation time and thus supporting the involvement of the water bridge in maintaining the interactions between AtENT3 and the tZR side chain (Figure 5F). To assess the stability of interactions among tZR, Trp34, Tyr61, and Asp129, we calculated the distributions of the distances between the interacting atom pairs (those visualized in Figure 5E) during the three 100 ns-long simulations. These distributions show that the distance between the hydrogen atom of the side-chain hydroxyl of tZR (H5) and the carboxylic oxygen of Asp129 (OD1), as well as the distance between H5 atom of tZR and the phenolic oxygen of Tyr61 (OH), oscillate around 4 Å. Assuming that the donor-acceptor distance in a typical hydrogen bond is less than 3.5 Å (Lemkul, 2019), the distribution of H5-OD1 and H5-OH distances supports the previous conclusion that the tZR-Asp129 and tZR-Tyr61 interactions are mediated by a water bridge, as the most likely distances are longer than the 3.5 Å threshold. The distribution of distances between the OD1 atom of Asp129 and the phenolic hydrogen (HH) of Tyr61 shows a sharp peak around 2 Å, indicating a stable hydrogen bond between these two atoms. This conclusion is also supported by the distribution of sizes of the angle formed by OD1, HH, and OH atoms, where the average angles are about 160°, i.e. close to a straight line (Figure S5). The distribution of distances between the oxygen of the side-chain hydroxyl of tZR (O5) and the nitrogen-bound hydrogen of Trp34 (HE1), ranging from 2 to 5 Å, does not show any significant peak, indicating that there are no stable interactions (Figure 5G). Distributions of distances involving the OD1 atom of Asp129 show a secondary peak, which is caused by the flipping of the Asp129 carboxyl group during the simulations. Altogether, the results from the docking and molecular dynamics show that Tyr61 and Asp129 of AtENT3 can stabilize AtENT3-tZR binding via interactions with the side-chain hydroxyl group of tZR, which might explain the preference of AtENT3 towards tZR over iPR (Figure 4C).

### 5.5 Loss of the AtENT3 Function Affects Shoot Development and *WUSCHEL* Expression in *A. thaliana*

To assess the physiological significance of the membrane transport of ribosylated CKs, we examined the phenotype of *A. thaliana* plants mutated in *atent3*, whose transport activity we have already characterized in this work. Given the previous report on the effects of *atent3* mutation on the primary root length (Korobova et al., 2021), we have focused on the plant shoots. We imaged shoots of 8-day-old wild-type *A. thaliana* plants and *anent3* mutants grown on the agar and 8, 11, and 15-day-old plants (wild type and *atent3*) grown on the cultivation substrate in pots. We processed the obtained image to measure the area of plant shoots from top view. The shoots of *atent3* plants are larger than the corresponding control in all cases (Figure 6A-B). For all measured parameters, see Table S15-16. For the statistical analysis of measured areas, see Table S17.

**Figure 6:**
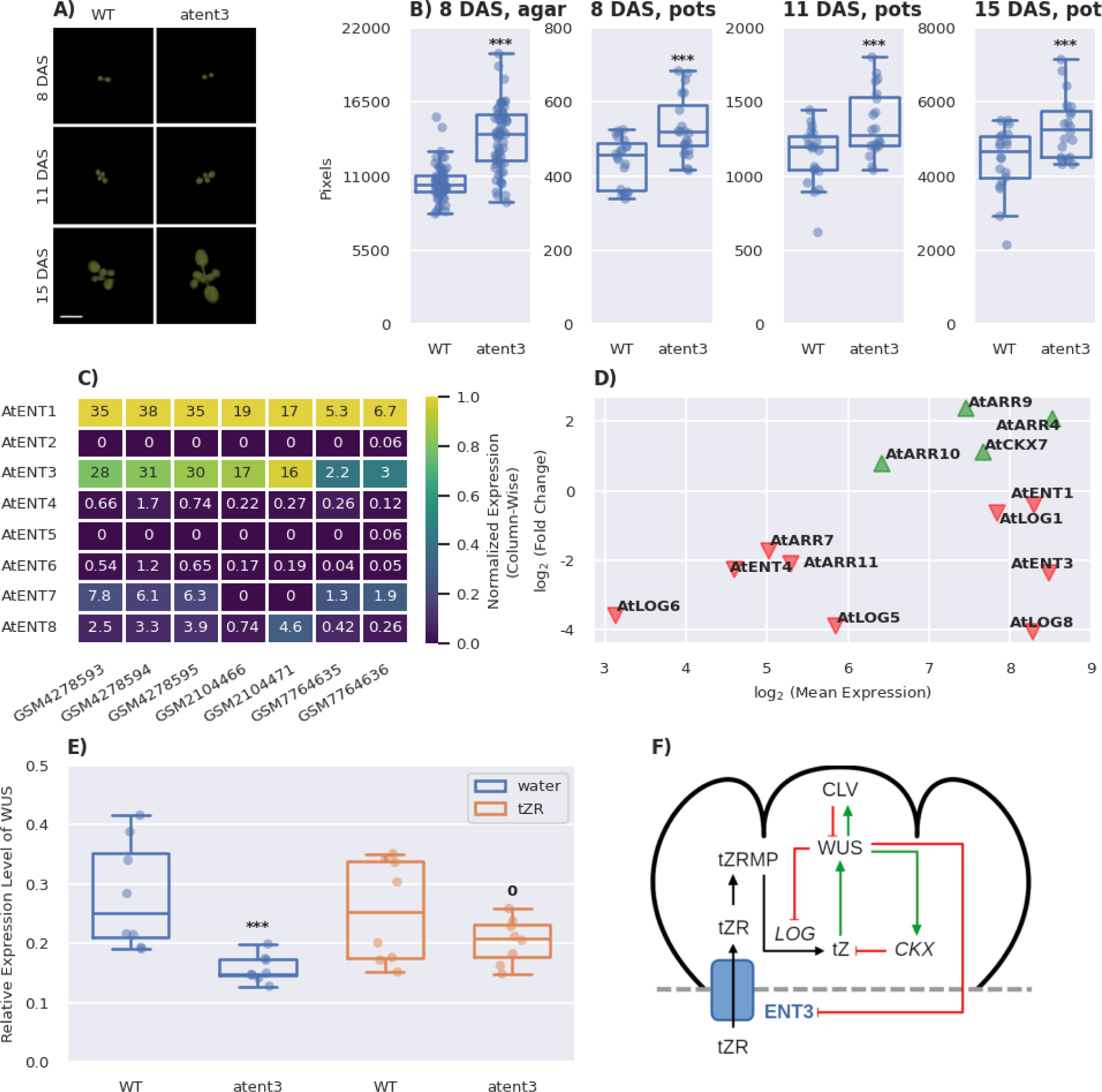
AtENT3-mediated transport of tZR contributes to the shoot development in *Arabidopsis thaliana*. **A**: Top view images of wild-type and *atent3 A. thaliana* plants grown in pots. Scale bar: 1 cm. **B**: Shoot areas of *A. thaliana* plants grown on the agar and in pots measured through image analysis. **C**: Relative expressions of *AtENT* genes in the shoot apices or apex-enriched tissues of *A. thaliana* retrieved from the Gene Expression Ominubs (GEO) database via accessions GSM4278593-95 (Yang et al., 2021), GSM2104466 and 71 (Mandel et al., 2016), and GSM7764635-36 (Incarbone et al., 2023). The colour scale is normalized from 0 to the maximal value in each column. **D**: Differential expression of genes related to CK transport, metabolism, and signalling in plants ectopically overexpressing *WUSCHEL* (*WUS*) in comparison to control plants. Data obtained from GEO accession GSE122610 (Ma et al., 2019). **E**: Relative expression levels of *WUS* in the shoots of 8 day-old agar-grown *A. thaliana* plants obtained through quantitative PCR. **F**: A schematic proposition of the function of AtENT3-mediated tZR transport in the maintenance of cytokinin homeostasis and WUS activity in the SAM. Black arrows denote movement and conversions of cytokinin species, green arrows activation, and red lines with flat ends inhibition. *P*-values obtained from the one-way ANOVA test comparing wild-type and *atent3* plants: 0 (*P* > 0.1), * (0.1 >= *P* > 0.05), ** (0.05 >= *P* > 0.01), *** (*P* <= 0.01). DAS: days after sowing, LOG: LONELY GUY, CKX: cytokinin dehydrogenase, CLV: CLAVATA.

The effect of the *atent3* mutation on the shoot development might be related to the previously reported requirement of tZR for proper modulation of physiological responses in the shoot apex (Landrein et al., 2018; Lopes et al., 2021), which implies the presence of CK riboside-recognizing transporters responsible for supplying root-borne tZR to the the apex (Sakakibara, 2021). In the following experiments, we therefore investigated the possibility that one of these tZR-providing transporters is AtENT3.

To determine whether *AtENT* genes are expressed in the apex area, we examined transcriptomic data obtained from isolated apices or apex-enriched tissues of *A. thaliana* deposited to the GEO database under accessions of GSM4278593-95 (Yang et al., 2021), GSM2104466 and 71 (Mandel et al., 2016), and GSM7764635-36 (Incarbone et al., 2023). From these data, we extracted expression levels of *AtENT1-8*. In all samples, *AtENT1* and *3* are abundantly expressed, sometimes followed by *AtENT7* and *8* (Figure 6C), supporting the idea that AtENT3 supplies the shoot apex with root-borne tZR.

As it was previously shown that increased CK supply leads to upregulation of *WUSCHEL* (*WUS*) expression in the shoot apical meristem (SAM) (Landrein et al., 2018), we further examined a possible relationship between the expression of *WUS* and *AtENT3*. We examined transcriptional data under accession GSE122610, where the authors evaluate the effects of ectopic *WUS* overexpression in 5-day-old *A. thaliana* seedlings (Ma et al., 2019). In this dataset, we searched for genes related to CK metabolism, transport or signalling. Overexpression of *WUS* leads to downregulation of the transporters *AtENT3* and *4*, CK-activating genes from the LONELY GUY (LOG) family, *AtLOG5*, *6* and *8*, and CK-responsive genes from the ARABIDOPSIS RESPONSE REGULATOR (ARR) family, *ARR7* and *11*. Conversely, *ARR4*, *9,* and the CK-degrading CYTOKININ DEHYDROGENASE 7 (*AtCKX7*) are upregulated (Figure 6D).

Having seen that overexpression of *WUS* can affect the expression of *AtENT3*, we next examined the expression of *WUS* in the shoots of *atent3* mutant via quantitative PCR. To determine whether potential changes in *WUS* expression are due to a lack of ribosylated CKs in the shoot apex, we treated half of the wild-type plants and *atent3* mutants with 1 µM tZR in water and the remaining plants with the corresponding amount of water alone. The expression of *WUS* is lower in the water-treated *atent3* mutant than in the corresponding wild type plants. Treating the plants with tZR partially reduces this difference in *WUS* expression (Figure 6E). For the relative expression levels of *WUS* and their statistical analysis, see Table S18 and Table S19, respectively. Both these findings support the hypothesis that AtENT3 provides the apex with root-borne tZR. Based on our findings, we have proposed an updated working scheme explaining the role of root-borne tZR in the shoot apex. This scheme, depicted in Figure 6F and further discussed below, will be the pivot focus of our future research.

## 6 Discussion

Ribosylated CKs are the dominant CK form transported through the xylem and phloem (Corbesier et al., 2003; Sakakibara, 2021; Takei et al., 2001) Their effective distribution between the cellular and extracellular compartments, mediated by membrane-bound carriers, is thus a crucial aspect of communication among different tissues and organs. In this work, we address particular differences between the transport kinetics of CK nucleobases and ribosides via radio-accumulation assays in BY-2 cell culture, a plant model system. We show that the uptake kinetics of CKs with isoprenoid chains differ significantly more between nucleobases and ribosides than among compounds with different side chain compositions, highlighting the presence of CK riboside-specific transporters (Figure 1). Conversely, the uptake kinetics of aromatic CKs (BA and BAR) do not differ, suggesting that aromatic CKs are recognized by a different transport system. The specific transport properties of BA and BAR evoke inquiries about the roles of aromatic CKs in plants and the means of maintaining their homeostasis in general.

We further show that the CK uptake occurs via at least two different systems of membrane-bound carriers. One of these systems exclusively recognizes ribosylated substrates. The other one primarily transports nucleobases but not as strictly as the first system. We derive this conclusion from the general trend of our data from accumulation assays, which shows that the inhibition of the CK riboside uptake by nucleobases is weaker than when the roles are reversed (Figure 3). The existence of riboside-specific transporters hypothetically allows plants to regulate the distribution of ribosylated CKs in a targeted manner. Since ribosylated CKs are primarily transported over long distances, the CK riboside-specific carriers could be found within or close to vascular tissues. This particular expression pattern could serve as a clue in the search for other CK riboside transporters in future research.

We used AtENT3 as a representative membrane-bound carrier of CK ribosides to further characterize the CK riboside transport. Our measurements of CK uptake in the *AtENT3*-expressing BY-2 cells showed that AtENT3 transports CK nucleobases and ribosides alike and thus does not belong to the CK riboside-specific carriers (Figure 4). Similar measurements focused on the ability of other AtENTs to distinguish between CK nucleobases and ribosides would allow us to tell whether this trend observed for AtENT3 applies to all AtENTs or whether the family includes other members, that may be specific for CK ribosides. The docked poses of tZ and iP (Figure S1 and S3) show that both nucleobases interact with the Gln62 residue of AtENT3 via a hydrogen bond. Gln62 is conserved among AtENT2-7, while in AtENT1 and 8, the corresponding position is occupied by methionine, an amino acid with an aliphatic side chain that is unlikely to form the mentioned hydrogen bond (Figure 5B). We propose that due to this difference in the amino acid composition, AtENT1 and 8 will not recognize CK nucleobases as substrates, which will be interesting to prove or disprove in future experiments.

The character of the position corresponding to Gln62 in AtENT3 also varies among other ENTs listed in Figure 5B. Notably, this position differs between HsENT1 (methionine) and HsENT2 (glutamine), cobsistent with their previously reported affinities towards nucleobases and nucleosides. HsENT1 favours uridine, a riboside, over nucleobases adenine, thymine, and hypoxanthine, whereas HsENT2 transports the nucleobases with affinities equal to that towards uridine or greater (Yao et al., 2011). The differential affinity towards nucleobase substrates between HsENT1 and 2 supports the hypothesis that the variable nature of the position corresponding to Gln62 in AtENT3 can affect the substrate specificities of ENTs. Another example is, PfENT1, which has the position Gln62 position occupied by glutamate and transports inosine, a riboside, and hypoxanthine with comparable affinities (Wang et al., 2023).

We have further observed that AtENT3 prefers tZR over iPR (Figure 4). The two substrates differ in the hydroxylation status of their side chains, as tZR is hydroxylated and iPR is not. Through the molecular docking of tZR in the predicted structural model of AtENT3 and subsequent molecular dynamic simulations of the AtENT3-tZR complex, we have identified stable interactions among residues Tyr61, Asp129 and the side-chain hydroxyl group of tZR. These interactions could explain the preference of AtENT3 for tZR. Both Tyr61 and Asp129 are conserved among all AtENTs but not among animal and other ENTs (Figure 5B), suggesting that their presence allows preferential binding of tZ-derived CKs by AtENTs. To investigate these indications further, it would be worth comparing affinities towards tZR and iPR for other AtENTs. as the preference towards the former should be conserved alongside Tyr61 and Asp129 residues. Another option to test the involvement of Tyr61 and Asp129 residues in stabilizing tZR is to measure the transport of tZR mediated by non-plant ENTs.

In the last part of this work, we have addressed the physiological impact of the AtENT3-mediated transport in *A. thaliana* shoots. The membrane transport of CK ribosides has been deemed a necessary step for the activation of tZR coming via xylem from roots up to the shoot apex and subsequent stimulation of *WUS* expression (Davière and Achard, 2017; Landrein et al., 2018; Lopes et al., 2021; Osugi et al., 2017; Sakakibara, 2021). This fact prompted us to investigate whether a change in *AtENT3* expression affects shoot development. Combining publicly available transcriptomic data with our quantitative PCR measurements has revealed that overexpression of *WUS* downregulates expression of *AtENT3* and that the *atent3* mutation downregulates expression of *WUS*. The shoots of *atent3* plants show larger cotyledons than the wild-type plants, which resembles the previously reported phenotype of *wus* mutant seedlings (Hamada et al., 2000; Laux et al., 1996). Nevertheless, this phenotype can also stem from the reduced retention of tZR in *atent3* roots (Korobova et al., 2021).

Other genes affected by *WUS* oversexpression include downregulated *ENT4*, *LOG5, LOG6*, *LOG8* (to a lesser extent also *ENT1* and *LOG1*) and upregulated *CKX7,* indicating that the overexpressed *WUS* tends to inhibit CK signalling at several levels as negative feedback. This feedback likely occurs via activation of type-A ARRs, which are typically characterized as CK-repressive (To et al., 2007, 2004). We have seen that overexpression of *WUS* upregulates type-A *ARR4* and *ARR9*. On the other hand, type-A *ARR7* is downregulated, suggesting that ARR7 does not participate in attenuating the incoming CK signal but rather in further regulation of the shoot apex development, consistent with a previous report (Leibfried et al., 2005). The differential effect of *WUS* overexpression on the type-A ARRs could also be a hint of plants being able to discern between CK signalling over long distances (which modulates *WUS* activity in response to environmental cues) and at the local level (which further shapes the SAM) by employing different response regulators for each. The suggested role of AtENT3 in the regulation of *WUS* expression and the subsequent WUS feedback are schematically depicted in Figure 6F, and its further assessment is another perspective of our future research.

## 2 List of Abbreviations

ABC: ATP-BINDING CASSETTE
ADFR: AutoDockFR software suite
ARR: ARABIDOPSIS RESPONSE REGULATOR
ANOVA: analysis of variance
At: mouse-ear cress (*Arabidopsis thaliana*)
AZG: AZA-GUANINE RESISTANT
BA: benzyladenine
BAR: benzyladenosine
Bt: cattle (*Bos taurus*)
BY-2: Bright Yellow 2
CCCP: carbonyl cyanide 3-chlorophenylhydrazone
CK: cytokinin
CKX: CYTOKININ DEHYDROGENASE
CMOS: complementary metal-oxide-semiconductor Col-0 Columbia-0
cZ: *cis*-zeatin
DHZ: dihydrozeatin
DHZR: dihydrozeatin riboside
DiPy: dipyridamole
DMSO: dimethyl sulfoxide
Dr: zebrafish (*Danio rerio*)
ENT: EQULIBRATIVE NUCLEOSIDE TRANSPORTER
Hs: human (*Homo sapiens*)
iP: isopentenyl adenine
iPR: isopentenyl adenosine
LOG: LONELY GUY
MAD: median of absolute deviation
Mm: mouse (*Mus musculus*)
MS: Murashige-Skoog
NBTI: *S*-(4-nitrobenzyl)-6-thioinosine
NCBI: National Center for Biotechnology Information
Os: rice (*Oryza sativa*)
Pf: *Plasmodium falciparum*
PUP: PURINE PERMEASE
REL: relative expression level
Rn: rat (*Ratus norvegicus*)
SAM: shoot apical meristem
Sc: yeast (*Saccharomyces cerevisiae*)
SWEET: SUGAR WILL EVENTUALLY BE EXPORTED TRANSPORTER
TM: transmembrane helix
WUS: WUSCHEL
tZ: *trans*-zeatin
tZR: *trans*-zeatin riboside

## 7 Author Contribution

DN, KH, MH, OP designed the experiment and conception. MH, KM performed molecular techniques. PK, DN, KH performed transport assays. PH, DN constructed mathematical model. JL, JS, MP, PK, VM performed phenotypical analysis. DN, VM, and KH wrote the manuscript. All authors read and approved the manuscript.

## 8 Acknowledgements

The authors wish to thank to Julie Talpová and Anita Bírošíková for excellent technical support and Martin Lepšík and Roman Pleskot for constructive remarks on computational methods.

## 10 Supplementary Files

**Table S1**: Sequences of primers used for cloning and quantitative PCR.

**Table S2**: Parameters for molecular dynamic simulations.

**Table S3**: Kinetic parameters obtained through the mathematical modelling of accumulations assays with radio-labelled tZ, tZR, iP, iPR, BA, BAR, DHZ, and DHZR in BY-2 cells.

**Table S4**: Statistical analysis of data from Table S3.

**Table S5**: Kinetic parameters obtained through the mathematical modelling of accumulation assays with radio-labelled tZ, tZR, iP, and iPR and their non-labelled counterparts at various concentrations as inhibitors.

**Table S6**: Kinetic parameters obtained through the mathematical modelling of accumulation assays with radio-labelled tZ, tZR, iP, and iPR with our without the addition of 10 µM CCCP at the beginning of the assay.

**Table S7**: Statistical analysis of data from Table S6.

**Table S8**: Kinetic parameters obtained through the mathematical modelling of accumulation assays with radio-labelled tZ, tZR, iP, and iPR with the addition of 10 µM CCCP at *t* = 420 s.

**Table S9**: Kinetic parameters obtained through the mathematical modelling of accumulation assays with radio-labelled tZ, tZR, iP, iPR, BA, and BAR and various nucleobases and ribosides at the concentration of 20 µM as competitors.

**Table S10**: Expression levels of tobacco *ENT* homologs in BY-2 cells. Data taken from GSE160438.

**Table S11**: Kinetic parameters obtained through the mathematical modelling of accumulation assays with radio-labelled tZ, tZR, iP, and iPR in control and induced BY-2 cells harbouring the *AtENT3* gene.

**Table S12**: Statistical analysis of data from Table S11.

**Table S13**: Kinetic parameters obtained through the mathematical modelling of accumulation assays with radio-labelled tZR and NBTI, Dipy or CCCP as inhibitors in control and induced BY-2 cells harbouring the *AtENT3* gene.

**Table S14**: Statistical analysis of data from Table S13.

**Table S15**: Parameters obtained through processing of images of 8-day-old wild-type and *atent3 A. thaliana* plants grown on the agar.

**Table S16**: Parameters obtained through processing of images of 8, 11, and 15-day-old wild-type and *atent3 A. thaliana* plants grown in pots.

**Table S17**: Statistical analysis of areas from Table S16 and Table S17.

**Table S18**: Relative expression levels of *AtWUS* measured through quantitative PCR in the shoots of 8-day-old wild-type and *atent3 A. thaliana* plants treated with 1 µM tZR in water or the corresponding amount of water.

**Table S19**: Statistical analysis of data from Table S18.

**Supplementary methods**: Derivation of models given by equation (1) and equation (2).

**Figure S1-S4**: The best-docked positions of tZ, tZR, iP, and iPR in the AlphFold-predicted structure of AtENT3.

**Figure S5**: Distribution of distances and angles during molecular dynamic simulations of the AtENT3-tZR complex.

